# OmicsTIDE: Interactive Exploration of Trends in Multi-Omics Data

**DOI:** 10.1101/2021.02.01.428836

**Authors:** Theresa Harbig, Julian Fratte, Michael Krone, Kay Nieselt

**Affiliations:** Institute for Bioinformatics and Medical Informatics, University of Tuebingen, Tuebingen, 72076, Germany

## Abstract

**Motivation:** The increasing amount of data produced by omics technologies has significantly improved the understanding of how biological information is transferred across different omics layers and to which extent it is involved in the manifestation of a given phenotype. Besides data-driven analysis strategies, interactive visualization tools have been developed to make the analysis in the multi-omics field more transparent. However, most state-of-the-art tools do not reconstruct the impact of a given omics layer on the final integration result. In general, the amount of omics data analyses strategies and the fields of applications lack a clearer classification of the different approaches.

**Results:** We developed a classification for omics data focusing on different aspects of multi-omics data sets, such as data type and experimental design. Based on this classification we developed the Omics Trend-comparing Interactive Data Explorer (`OmicsTIDE`), an interactive visualization tool developed to address the limitations of current visualization approaches in the multi-omics field. The tool consists of an automated part that clusters omics data to determine trends and an interactive visualization. The trends are visualized as profile plots and are connected by a Sankey diagram that allows an interactive pairwise trend comparison to discover concordant and discordant trends. Moreover, large-scale omics data sets are broken down into small subsets of concordant and discordant regulatory trends within few analysis steps. We demonstrate the interactive analysis using `OmicsTIDE` with two case studies focusing on different types of experimental designs.

**Availability:** `OmicsTIDE` is a web tool and available via http://tuevis.informatik.uni-tuebingen.de/

## 1 Introduction

With the advent of *high-throughput technologies*, it has become affordable to comprehensively study the flow of information by taking into account the entirety of all genes, transcripts, proteins, or metabolites within a sample (Hasin *et al*., 2017). A common approach is to analyze a single omics layer, such as transcriptomics, proteomics, or metabolomics, under different experimental conditions. On the other hand, the analysis of different omics layers under one specific experimental condition is applied to focus on how the flow of biological information propagates through different layers. This approach is for instance used to determine correlations between the abundances of expressed transcripts and translated proteins in a sample.

With the increasing complexity and amount of data deriving from high-throughput technologies, such as next-generation sequencing or mass-spectrometry-based approaches, that include multiple omics layers, the demand for methods that integrate the different omics data sets has been steadily increasing over the past decades (Subramanian *et al*., 2020).

While a data-driven integration can derive interesting coherences between different omics layers from the data, this process often remains intransparent. For instance the impact of single genes or groups of genes on the integration is often not easy to determine. To overcome this limitation of the purely data-driven approaches and to show how the single components of an omics-integration study contribute to detect concordant and discordant patterns in the compared abundance of omics data, different approaches have been developed (Hernández-de Diego *et al*., 2018; Vercruysse *et al*., 2020). Many of these approaches reduce the complexity of the data sets by classifying single genes into different categories based on their “behaviour”. For example, in a data set that deals with only two conditions, a gene could be classified as being *up-regulated* in one condition with respect to the the other condition. The situation becomes more complex if more than two conditions or even a time series are given in a data set. This requires the application of clustering methods to obtain representative *trends* for sets of genes. Here, we define a *trend* in omics abundance data as a set of elements (such as genes, proteins, or metabolites) that follow a distinct course across more than two conditions.

In order to provide a tool that overcomes the current limitations in the omics visualization field, we first sought to devise a general classification system for omics data. This classification builds the framework for the *Omics Trend-comparing Interactive Data Explorer* (`OmicsTIDE`), a tool that creates a connection between the single genes and the trends derived from multi-omics data sets.

`OmicsTIDE` visualizes trends as profile plots, also known as parallel coordinate plots (Heinrich and Weiskopf, 2013), and adapts the idea of summarizing and visualizing similarities and differences in two data sets by using a Sankey diagram (Lex *et al*., 2012). The Sankey diagram is used to compare the trends across the two data sets, i.e., the genes following either concordant or discordant trends. `OmicsTIDE` embeds trend comparison in an analysis that breaks down large-scale data sets into small subsets within a few steps based on selecting groups of genes in the Sankey diagram. By additionally allowing several pairwise comparisons within one single analysis, `OmicsTIDE` aims to combine the insights from different pairwise comparisons into one large analysis. We demonstrate the effectiveness of `OmicsTIDE` in two case studies dealing with different experimental designs.

## 2 Related Work

The integration of multi-omics data has become a steadily growing research field. For this paper we define a multi-omics tool as a tool that integrates and visualizes data of two or more omics layers in a parallel fashion. In this related work section, we thereby focus on tools that analyze different omics data in a combined instead of a separated or sequential manner.

The most straightforward way of visually analyzing multi-omics data is mapping them directly to a genome sequence or a pathway. Any kind of omics data that can be mapped to a genome sequence can be represented in genome coordinate-based visualizations such as genome browsers (Nusrat *et al*., 2019). With tracks stacked upon each other, various omics layers can be displayed simultaneously. Another common approach is mapping omics data to a pathway ID in a node-link diagram, where genes, proteins, and metabolites can be shown simultaneously (Luo and Brouwer, 2013; Eichner *et al*., 2014). While genome browsers and pathway maps can comprehensively and intuitively visualize multi-omics data, they usually do not aim at visualizing the entirety of the data but only a small window of the genome or a single pathway of interest that can be determined using, for example, pathway enrichment methods (Huang *et al*., 2009; Eichner *et al*., 2014). Additionally, when displaying omics data in multiple conditions, both types of visualization can become hard to interpret due to overplotting.

Besides these straightforward visualization techniques, various computational methods for the integration of multi-omics data have been developed. These approaches make use of different techniques, including network-based methods (Yan *et al*., 2018), matrix factorization (Huang *et al*., 2017), or Bayesian methods (Bersanelli *et al*., 2016). Often omics data are clustered to obtain trends, which can be as simple as classifying abundance data in time series experiments as decreasing or increasing over time in a two condition experiment (Hackett *et al*., 2015). If more conditions are given in a data set the number possible trends increases, which often requires more advanced clustering approaches (Rappoport and Shamir, 2018; Tini *et al*., 2019). These approaches can be divided into *early integration* and *late integration* approaches. While early integration approaches first concatenate the data of the different omics layers and then cluster the merged data, late integration methods first find patterns in the features of each layer separately which can be combined as input for a regression or classification (Sharifi-Noghabi *et al*., 2019).

Commonly, the results of the integration methods are visualized in node-link diagrams or in trend visualizations, such as heatmaps and profile plots. `3Omics` is a tool for the integration and visualization of multi-omics data, which integrates up to three different omics layers, e.g. transcriptomics, proteomics, and metabolomics data. `3Omics` uses hierarchical clustering and visualizes the results as a clustered heatmap. Alternatively it creates correlation networks as node-link diagrams. For network visualizations the same limitations arise as for pathway visualizations, since with an increasing number of nodes, the single connections cannot be perceived well. A similar heatmap visualization as provided in `3Omics` has been implemented in the recently published tool `multiSLIDE`, which combines two heatmaps side-by-side comparing transcriptomics and proteomics data (Ghosh *et al*., 2020). While heatmaps provide an intuitive way of visualizing abundance data, they can become huge when analyzing a large number of genes. Due to their size and because of the usage of the color encoding, trends may become difficult to determine (Gehlenborg, 2012).

`Paintomics` follows an alternative approach to integrate multiple omics levels and to include large-scale data sets in the analysis (Hernández-de Diego *et al*., 2018). First, it combines the data from the different omics experiments by determining the `KEGG` pathways in which the genes can be found. This allows the integration of a large number of different omics layers by finding a common pathway key for a shared integrated analyses. Based on the determined pathway, different analyses can be performed, such as pathway enrichment or other pathway network analyses. For the results of the pathway analysis, trends of the given omics layers can be analyzed by extracting the information of how the genes of an omics layer are reflected in a selected pathway. However, it does not show to what degree the genes within a given data set contribute to the final trend. To directly address the question of how the similarities and differences of trends between two data sets can be used to provide information about underlying mechanisms, different strategies to compare trends have been developed.

For instance, an approach to visualize and compare trends was demonstrated in a recently published study on the comparison of the transcriptomes of *Arabidopsis thaliana* and *Zea mays* (Vercruysse *et al*., 2020), where trends in orthologous genes in leaf development were determined and compared. For the visualisation of trends, the authors use profile plots. Thereby trends can be determined more easily than it would be possible in a heatmap. The approach comprises the determination of trends by hierarchical clustering and the subsequent comparison of these trends. Although this comparative approach using hierarchical clustering provides a good overview of the trends in the two data sets, it has several limitations. First of all, the hierarchical clustering requires to first perform a manual separation of the hierarchical clustering into sub-clusters to categorize the single trends which might be a tedious and error-prone work. Secondly, the connection between the single trends for either being concordant or discordant has to be guided by statistical measures. Finally, the analysis and visualization is static and does not allow user interactions to extend the analysis with users knowledge.

Despite the fact that many tools have been developed to extract common features from integrated omics data, interactive visualization tools that make the data integration transparent and interpretable are rare. As shown exemplarily for `3Omics` and `Paintomics`, the major drawback of the current state-of-the-art tools is that they do not create a connection between the genes in the single data set and the “global” trends derived by integrating the different data sets. This limitation might be due to the fact that the resolution for single genes decreases with the growing number of integrated data sets. Hence, creating a connection between the single links and their trend is crucial to overcome this limitation. This was the main motivation for the development of `OmicsTIDE`. Before we introduce the design of `OmicsTIDE` we will first turn to the question of how to conceptually classify omics data. This classification builds the requirement framework for `OmicsTIDE`.

## 3 Classification of Omics Data

In order to create an abstract representation of omics data, which helps us to further identify the requirements for a novel multi-omics tool from a visualization point of view, we developed a classification of omics data. First, omics data can be classified by the *attribute type* (Figure 1a). Data that is related to genomics research, typically comes with categorical data, e.g. for SNP analysis, which is usually provided as a *Variant Call Format* (`VCF`) file. In contrast, the majority of omics technologies, such as transcriptomics, proteomics, or metabolomics deals with quantitative data. Quantitative data (or abundance data) is usually provided in a data matrix, with *n* rows, corresponding to the genes for example, and *p* columns, corresponding to conditions (where usually *n ≫ p*). Additionally, the abundance data contains a key attribute, such as a gene ID that uniquely defines each row. Abundance matrices are the common data format that can be easily processed and combined with other matrices sharing the same keys.

**Fig 1:**
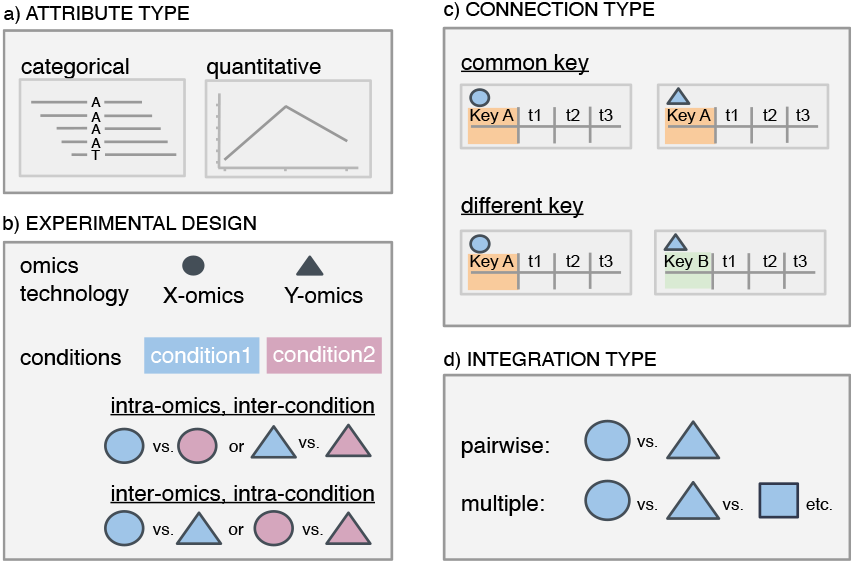
Omics data can be classified in different ways that is presented here for four categories exemplarily. (a) The output of omics experiments can be described as a matrix of genes and conditions.The *attributes* are either of categorical (mutations) or quantitative (e.g. transcript/protein levels) type. (b) Omics data analysis can also be classified by the *experimental design*: here, the analysis typically includes either the comparison of different conditions in the same omics layers (*inter-condition* and *intra-omics*) or the comparison of data from different omics layers within the same condition (*intra-condition* and *inter-omics*). (c) The combined analysis of omics data sets can be classified by whether the data sets can be joined by a common *key attribute* or not. (d) The *integration* of different omics layers can either be done in a pairwise fashion or by directly comparing multiple layers at once.

Secondly, comparative omics experiments can be classified by their experimental design, depending on the question one wants to pursue (Figure 1b). Abundance data usually contains the result of an experiment that was conducted within a given omics layer (*intra-omics*) and between different conditions (*inter-condition*). A biological question linked to this experimental setup could be for example “how does the transcriptome differ between a wild-type organism and an organism carrying a given mutation?”. Alternatively, omics experiments can answer additional biological questions when they are planned to include different omics layers (*inter-omics*) studying the same biological condition (*intra-condition*). Such experiments typically combine two or more different data modalities. In this case, the main interest may be to check how the flow of information changes between the different omics layers, e.g. between the transcriptome and proteome.

If the *inter-omics* approach is chosen as experimental design, the connection of integrated data sets can be created based on common key IDs with which the data sets can be combined or compared (Figure 1c). If the data sets do not share key IDs, a direct comparison cannot be conducted. An alternative approach to combine two data sets that do not share the same IDs could be to use meta-information, such as pathways IDs to make the two data sets comparable.

For *inter-omics* experimental designs, the decision on the number of omics layers that are used for the data integration determines the subsequent downstream analysis steps (Figure 1d). In order to study a given biological question it might be sufficient to compare two omics layers. More complex questions might require different omics studies (*multi-omics*). This approach has the advantage of a more powerful analysis to find specific patterns in the integrated data sets.

The choice for a suitable analysis approach always depends on the data as well as on the biological question one pursues. The overall aim for all multi-omics analyses is to perform a combined and not a sequential analysis and to derive patterns from the studied data. Integration methods often put a lot of emphasis on integrating as much information and as many layers as possible by applying sophisticated statistical methods. However, this often results in information of a specific integrated layer not being accessible to users. Furthermore, the integration of multiple layers often exceeds the scientific scope and budget of the majority of research groups in the life sciences. Hence, a simple tool is demanded that allows users to conduct straightforward visual-interactive analysis for both, *intra-* and *inter-omics* data sets. The main challenge in creating such a tool is providing an overview of the integration, while still showing information about single genes. With `OmicsTIDE` this challenge was addressed within the framework of the developed classification scheme.

## 4 Design and Implementation

Based on the classification system for omics data described in the previous section, we developed `OmicsTIDE`. Here, we describe the requirements and design decisions that were guided by this classification scheme. `OmicsTIDE` is an interactive *inter-omics* and *intra-omics*, as well as *intra-condition* and *inter-condition* visualization tool to compare trends of omics abundance data in a pairwise manner. The approach derives and visualizes trends for two-dimensional experimental designs. The first dimension is represented by the data sets that are compared, where each may be from one or two different omics layers. The other dimension is represented by further conditions that need to be consistent across data sets, such as the same time points or the same environmental conditions. Transcriptomic and proteomic data are especially suited for `OmicsTIDE` since they share common keys (gene IDs) and have quantitative attribute types. As transcriptomic and proteomic data can both be thought of as matrices showing abundance levels of genes at different conditions, they are identical from a data type perspective. This leads to a great flexibility in `OmicsTIDE` as a tool that can perform *inter-omics, intra-condition* comparisons is also able to perform *intra-omics, inter-condition* comparisons, such as comparing two transcriptomic data sets.

The central idea of our visualization approach is to cluster omics data sets containing abundance data and shared keys into trends that are visually connected via a Sankey diagram, which is a special kind of flow diagram showing the flow from one set of values to another. The trends of the different data sets are visualized adjacent to the *nodes* of the Sankey diagram. The height of the nodes encodes for the number of genes found in the trends, whereas the thickness of the bands (*links*) between the nodes encodes for the number of genes that either share trends in the two data sets (concordant genes) or show different trends (discordant genes).

The tool derives and visualizes the trend comparison by performing an analysis in three major steps referred to as *comparison selection, first-level analysis* and *second-level analysis* (Figure 2). The separation of the analysis is reflected in the dynamic tab-based design of `OmicsTIDE`, which guides the user through the analysis by gradually allowing to add new tabs corresponding to the respective analysis steps. This design facilitates to review, refine or even remove choices made in any tab in order to customize the analysis. The information gained by the user in this three-step analysis contributes to a more comprehensive understanding of the omics data.

**Fig 2:**
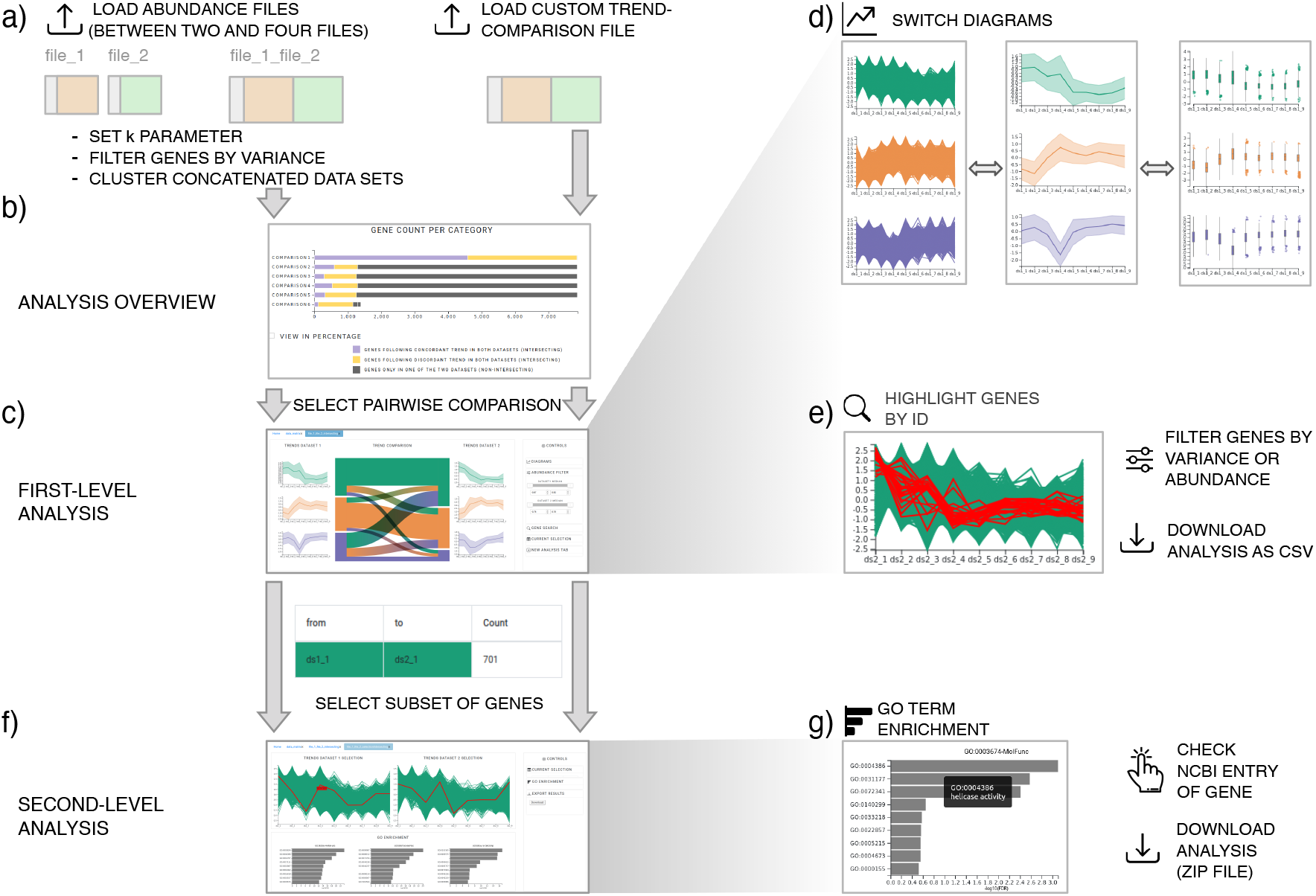
Basic workflow using `OmicsTIDE`. (a) Either multiple abundance files or a single custom trend-comparison file can be uploaded. (b) An overview of all conducted pairwise trend comparisons is shown as horizontal stacked bar chart showing the count of genes either being found in both compared files (intersecting) or only in one of the two files (non-intersecting). The number of intersecting genes in the bar chart is further categorized by either following a concordant (purple bar) or discordant trend (yellow bar) in the two compared files (c) After selecting a pairwise comparison in the overview visualization the data can be analyzed in the first-level analysis, consisting of a Sankey diagram comparing trends in both abundance files. (d) Users can switch between different trend diagrams, (e) highlight genes using gene ids and filter data by abundance and variance. (f) For a more detailed analysis subsets of genes can be analyzed in the second-level analysis which includes (g) viewing the NCBI entries of single genes and GO term enrichment analysis in a bar chart. The *x* axis corresponds to the *-log*_10_(*FDR*) values and the *y* axis corresponds to the ten most significant GO term hits.

### 4.1 Data Loading and Comparison Selection

The data input in `OmicsTIDE` offers two distinct data input options (Figure 2a). By choosing the default option, the user can load two or more abundance files that are clustered by `OmicsTIDE` to obtain trends. Choosing the second option, users can upload their own clustering results for a pairwise trend comparison created with any clustering algorithm.

After uploading abundance files, the input can be modified as follows. First, to restrict the analysis to variant genes, users can reduce the loaded data sets by removing genes based on the percentile range of their variances across different conditions within the respective data set. This prevents the formation of entirely flat trends or that the shape of the trend is influenced by low-variance genes. Next, to make the ranges of the data sets comparable, *z-score* normalization is applied. The crucial step for the trend comparison is the determination of the number of trends to be derived from the data. This is done by initializing the number of trends derived by k-Means clustering in the range between 2 and 6. For the clustering, `k-Means++` (Arthur and Vassilvitskii, 2006) is used. The fast clustering using k-Means allows users to quickly explore their data using different choices of k. For each pairwise combination, `OmicsTIDE` conducts two separate trend comparisons: One for the genes found in both data sets (intersecting genes) and one for the genes found only in one of the two data sets (non-intersecting genes). Here, `OmicsTIDE` makes use of an *early integration* approach (Rappoport and Shamir, 2018) by first concatenating the two data sets (e.g. for intersecting genes) in one combined matrix and applying the clustering to the combined matrix. With this approach, the genes can easily be classified into following concordant or discordant trends in the two data sets compared, which represents an intuitive concept for users. Such a concept would be harder to realize using *late integration* approaches which would produce different trends in both data sets.

An overview visualization of the clustering results helps users to select the comparison of interest for the trend exploration in the first-level analysis (Figure 2b). The comparisons are visualized using stacked bar charts showing concordant, discordant and non-intersecting genes. Together with the user’s knowledge about the experimental design, these three attributes are especially useful for selecting a comparison of interest since they represent the similarity of two data sets using very few values. Stacked bar charts are a simple yet effective visualization for this type of data, that can easily be interpreted. Moreover, to provide a first glimpse on the computed trends a small preview of the trend exploration visualization is shown, which is identical to the visualization in the first-level analysis.

### 4.2 First-level Analysis: Trend Exploration

The first-level analysis tab provides a detailed visualization of the selected trend comparison in a Sankey diagram (Figure 2c). The, trends, nodes and links are colored using a set of categorical colors. Commonly, the links in Sankey diagrams follow a gradient, transitioning from the color of the left node to the right node. Although this is often perceived as visually appealing, we decided to inverse this gradient to easily identify the set of trends in one data set connected to a trend in the other data set.

By default, the trends are visualized using *centroid profile plots*, which provide an overview of the trends by showing the centroid line as well as the standard deviation of the trend as a band. Alternatively, users can choose profile or box plots for visualizing the trends (Figure 2d). Profile plots provide a more detailed view on the composition of each trend. However, studying the single-gene level is more suitable for a low number of genes since the visualization of a high number of gene profiles results in overplotting. As a third option, the user can study the variation within a given trend in more detail by using box plots. In addition to the analysis of intersecting genes, the trend visualizations are used for analyzing non-intersecting genes as well. Since non-intersecting genes do not share identifiers, they are not connected with a Sankey diagram.

The nodes and links in the Sankey diagram can be hovered to study the single trends between the two data sets in more detail. Hovering over a node or link will set the focus to the hovered element and the corresponding detail diagrams. If a node is hovered, all connected links are included in the focus, while all other elements are reduced in their opacity (*focus-on-hover* strategy). Hovering given elements in the visualization updates the detail diagrams accordingly. This update is facilitated via an animated transition to allow the user to visually link the hovering and the data update.

Users can check their own hypotheses about gene sets of interest, such as pathways, and analyze their behaviour across trends and data sets. To include own knowledge into the analysis, genes of interest can be highlighted based on their gene IDs (Figure 2e). Users can directly type in one or more gene IDs into a text field or upload a text file with gene IDs. The genes in the diagram corresponding to the given IDs in the query are marked in red.

The concordance of trends might depend on different attributes. Typically, data sets are filtered by a given variance or (median) abundance range of the genes. For example, users might want to analyze if the trends show more concordance when only highly variant of highly abundant genes are included in the analysis. Therefore, `OmicsTIDE` supports range sliders to dynamically filter data by the percentile ranges of the variance or the median abundance of the genes (Figure 2c, right). The variance and the median abundance and their respective percentiles are calculated prior to z-score normalization. The variance filtering in the first-level analysis can be applied as an alternative or in addition of the variance filtering provided when loading the data. In contrast to the variance filtering before loading the data, which is considered a prepossessing step, the filtering in the first-level analysis allows users to explore different ranges of variances quickly.

### 4.3 Second-level Analysis: Detailed Trend Analysis

Sets of genes corresponding to trends or the intersection of trends can be analyzed in a more detailed analysis to find, for example, enriched functions. `OmicsTIDE` allows users to either select links or nodes in the visualization in order to extract a single subset or all subsets corresponding to a given node. To show an overview of the selection, a selection summary table placed in the controls side bar shows the source node and the target node of each selected link as well as the number of the corresponding genes (Figure 2c, lower panel). Thereby users can more easily compare the actual numbers of genes corresponding to a link.

Selected genes can then be analyzed in detail in the second-level analysis (Figure 2f). Users can study gene sets on the single-gene level by hovering the single gene profiles and accessing information of an individual gene by clicking and being redirected to the corresponding `NCBI` entry. Furthermore, the biological function of the selected gene subset can be studied by summarizing the results in a Gene Ontology context (Figure 2g). Thereby, users can find functions that are enriched in sets of genes that show a different or the same trend in the compared data sets and form hypotheses about the biological processes causing the patterns. The `PantherDB` API is used to perform a GO enrichment study on a given gene selection (Mi *et al*., 2019). The species corresponding to the analysis needs to be selected in the controls side bar. The GO enrichment analysis is limited to those species currently provided by `Panther`. The enrichment analysis is performed for the three main GO categories *molecular function* (GO:0003674), *biological process* (GO:0008150) and *cellular component* (GO:0005575) using *Fisher’s exact test* and a multiple test correction with *False Discovery Rate (FDR)*. The results are visualized as horizontal bar charts showing the negative logarithm of the *FDR*. Thereby users can quickly identify the most significant results. Hovering the single bars will show a tool tip with more detailed information on the given GO term (Figure 2d, right).

### 4.4 File Export

Results of the analysis with `OmicsTIDE` can be exported in several ways. The user can download the currently studied pairwise trend comparison as *CSV* file. This enables the user to load the pairwise comparison for later analysis without the need to repeat the steps until this point. Moreover, analysis results of the detailed trend analysis can be exported as a *ZIP* file that contains the two profile plots as *SVG* files, the current selection, and the results of the GO enrichment as *CSV* files.

### 4.5 Implementation

`OmicsTIDE` is a web-based client-server application that makes use of established `Python` data analysis libraries in the back-end to ensure a seamless communication with the front-end via the *web server gateway interface* (WSGI)-based framework `Flask` (Grinberg, 2018). This communication allows `OmicsTIDE` an efficient outsourcing of more complex computations, such as the trend determination via clustering, to the back-end. In contrast, smaller computations, such as data filtering, are performed directly in the front-end. The `JavaScript` library `D3.js` is used for creating interactive `SVG`-based visualizations (Bostock *et al*., 2011). The `jQuery` library (Severance, 2015) is used to modify specific `HTML` elements and to send requests to the back-end. To realize the dynamic tab-based interface that allows a straightforward workflow with activated and faded single tabs, the Bootstrap framework (Spurlock, 2013) is applied. The source code of `OmicsTIDE` is available at https://github.com/Integrative-Transcriptomics/OmicsTIDE.

## 5 Case Studies

To demonstrate the usability of the pairwise trend comparison approach in `OmicsTIDE`, we applied it to different combinations of omics data sets. Here, we demonstrate the usage of `OmicsTIDE` in two case studies. In the first case study, we show how the combined analysis of transcriptomics and proteomics data derived from different stages of human blood cells can be used to extract biologically relevant concordant as well as discordant trends with few clicks only. The second case study combines two pairwise trend comparisons to extract information from both, different experimental conditions and omics layers to demonstrate the synergy effect that can be achieved by `OmicsTIDE`.

### 5.1 Blood Cell Differentiation in Bone Marrow

Neutrophils are an essential part of the human immune system. They are differentiated in the bone marrow and released to the bloodstream. The regulation of the neutrophil differentiation is subject of the first case study, examining the so-called granulopoiesis *in vivo* (Hoogendijk *et al*., 2019). The experimental design uses both transcriptome and proteome data from five differentiation stages to find concordant and discordant trends across the two omics layers. The five stages are (pro)myelocytes (PMs), metamyelocytes (MMs), immature neutrophils with band-shaped nucleus (BN), mature neutrophils with segmented nucleus (SNs) and finally the actual peripheral mature neutrophils derived from the blood stream (PMNs). Here, we show how `OmicsTIDE` allows the user to easily explore the trends across the two examined omics layers by reproducing the findings made by Hoogendijk et al. in their study. The data was taken from the supplementary material of the publication that contained quantified transcripts and proteins in the form of FPKM and imputed *log*_2_ LFQ measures, respectively. For each of the five conditions, the data for three and four biological replicates were given for transcripts and proteins, respectively. The analysis was performed on the mean values of all biological replicates for each of the five conditions.

To explore the trends shown by the transcriptome and the proteome of different blood cell types, the selection of *k* = 4 initial clusters resulted in clearly distinguishable trends that are shown as centroid profile diagrams for either data set (Figure 3). Four of the main trend comparisons between transcriptome and proteome described in the paper on the neutrophil differentiation could be reproduced by `OmicsTIDE`. The authors mainly focused on main combinations of trends in the proteome and transcriptome (so-called modules) and classified them based on GO enrichment and the enrichment of specific database entries. The patterns of the trend comparisons described in the paper could also be visually identified using `OmicsTIDE` by simply hovering the single links in the Sankey diagram (Figure 3) for the categories “RNA-binding proteins” (Figure 3a), “granule development” (Figure 3b), “biosynthesis & development” (Figure 3c), and “ROS machinery” (Figure 3d). The set of genes showing an increasing pattern in the transcriptome and a decreasing pattern in the proteome are mainly related to the RNA-binding proteins (Figure 3a) which could be confirmed in the GO term enrichment of the molecular function category using the second-level analysis tab.

**Fig 3:**
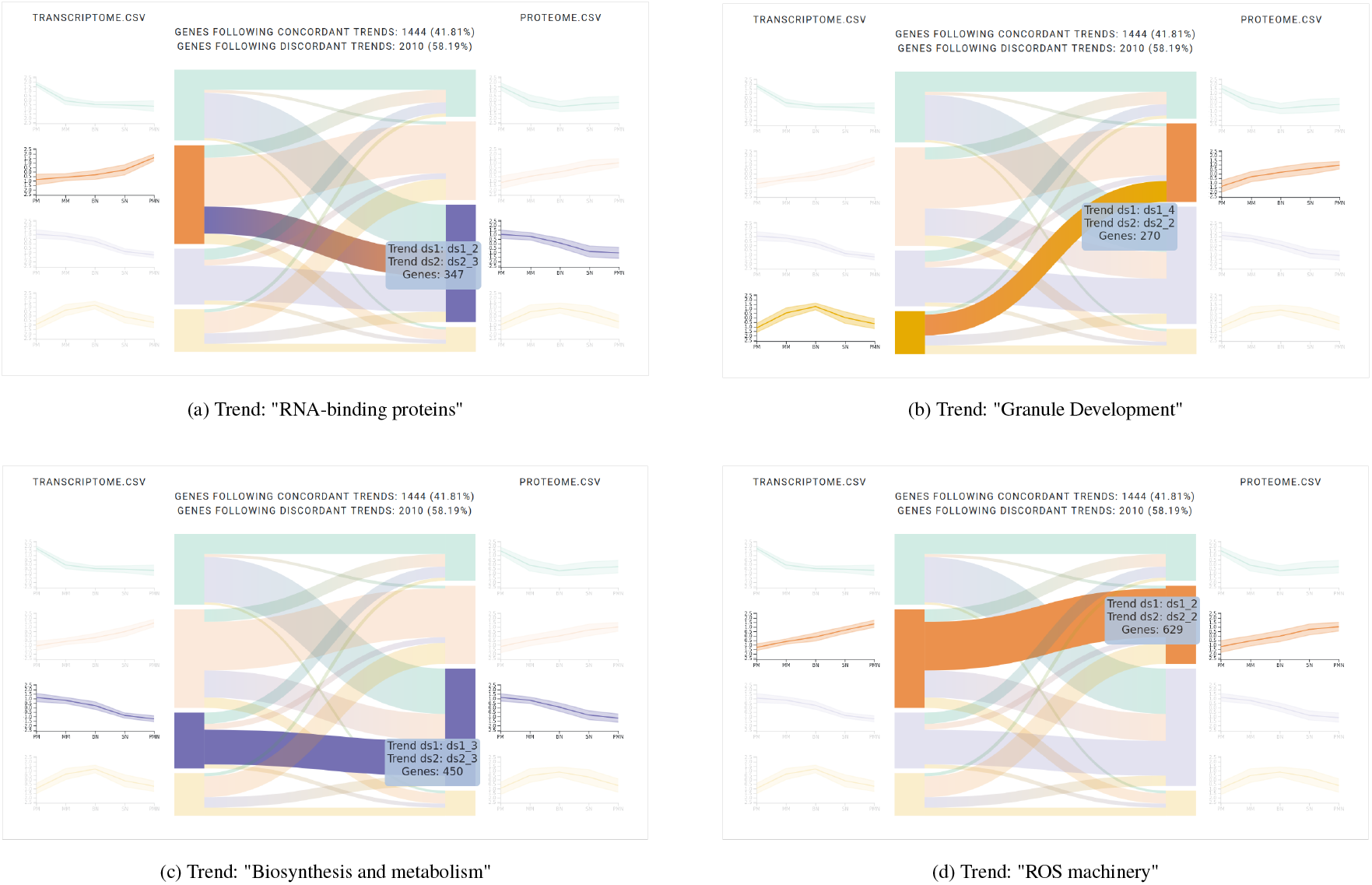
First-level analysis: Interactively studying the trend comparison between the transcriptome and proteome during blood cell differentation (Hoogendijk *et al*., 2019) by hovering reveals several concordant and discordant trends. (a-d) Four trends found by `OmicsTIDE` were associated with functionally annotated modules found in the study. The tooltip displays the information about the identity of the trends that are compared by the current hovering event and the number of contained genes.

Since the authors used a combination of GO enrichment and enrichment using other databases, the other functional annotations could not be easily reproduced. Yet, if we find that the combinations of trends in `OmicsTIDE` and the paper contain the same genes by manual comparison, we can conclude that the functional enrichment would show similar results as well. As the trends in `OmicsTIDE` and the modules in the paper are created differently, they are not directly comparable. However, the authors grouped the modules into different categories, such as concordant increasing, concordant decreasing, and increasing in the transcriptome while decreasing in the proteome. Moreover, in `OmicsTIDE` the sets of concordant genes stemming from both decreasing trends were merged. `OmicsTIDE` produced 629 increasing concordant genes, while 621 were found in the blood cell study with an overlap of 486 genes. Similarly, of the 1320 decreasing concordant genes, 1131 could be found in similar patterns in `OmicsTIDE` (total of 1439 trends). The other modules compared were much smaller and we found more genes in `OmicsTIDE`. Yet, we could find more than 70% of the genes of each module.

### 5.2 Transcriptome and Proteome Time Series Data Set of *Streptomyces coelicolor*

In order to demonstrate how *inter-omics* as well as *intra-omics* analysis can be combined using `OmicsTIDE`, we re-analyzed the data sets of a study exploring two *Streptomyces coelicolor* strains with respect to changes in their metabolisms under phosphate-starving growth conditions in a time-course experiment. The *Streptomyces coelicolor* strains M145 and M1152 were used to study the role of *biosynthetic gene clusters* (BGCs) for the production of antibiotics. M1152 is a genetically-engineered derivate of the M145 wild-type strain that was subject to the deletion of different BGCs (Gomez-Escribano and Bibb, 2011). For both strains samples were taken at several timepoints. Phosphate was depleted between timepoint 3 and timepoint 4 (Sulheim *et al*., 2020). Transcriptomics as well as proteomics data were produced across eight corresponding time points and for both strains. For each of the time points, three biological replicates were generated. Both, transcriptome and proteome data was initially quantified and *log*_2_-transformed. The data was normalized by an *intra-strain* and *intra-omics* quantile-normalization across all replicates. Finally, the mean of the three replicates was calculated for each strain, time point and omics layer separately.

In `OmicsTIDE` the four data sets (M145 transcriptome, M1152 transcriptome, M145 proteome, M1152 proteome) were loaded resulting in six pairwise trend comparisons. For the k-Means clustering *k* = 4 was chosen since it produced the most clearly distinguishable trends. In this application we first concentrated on the comparison of two different strains across a single omics layer (here M1152 transcriptome vs. M145 transcriptome). This approach points to find differences on the transcript level that reveal regulation differences between the wildtype and the mutant strain. The insights from this first pairwise comparison is then used to study whether and how these insights are also reflected by comparing the transcriptome and proteome within the mutant strain.

#### 5.2.1 *Intra-Omics*: M1152 transcriptome vs. M145 transcriptome

The comparison of the M1145 transcriptome and the M145 transcriptome revealed a total of 7,904 genes that appear in both data sets, whereof around 55% follow concordant trends (data not shown). After applying the abundance filtering to focus on the genes with a high median abundance of above the 80th percentile in both data sets the shape of the trends becomes clearly visible (Figure 4a). Interestingly, the green trend and the orange trend in the centroid profile plot show the exact inverse trend in the M1152 transcriptome. The same could be observed for the purple trend and the yellow trend. The inverse behaviour of the trends is also partly reflected in the M145 transcriptome. However, at least for these highly expressed genes, about 65% of the genes show discordant expression trends, pointing to also very different expression regulation in these two strains.

**Fig 4:**
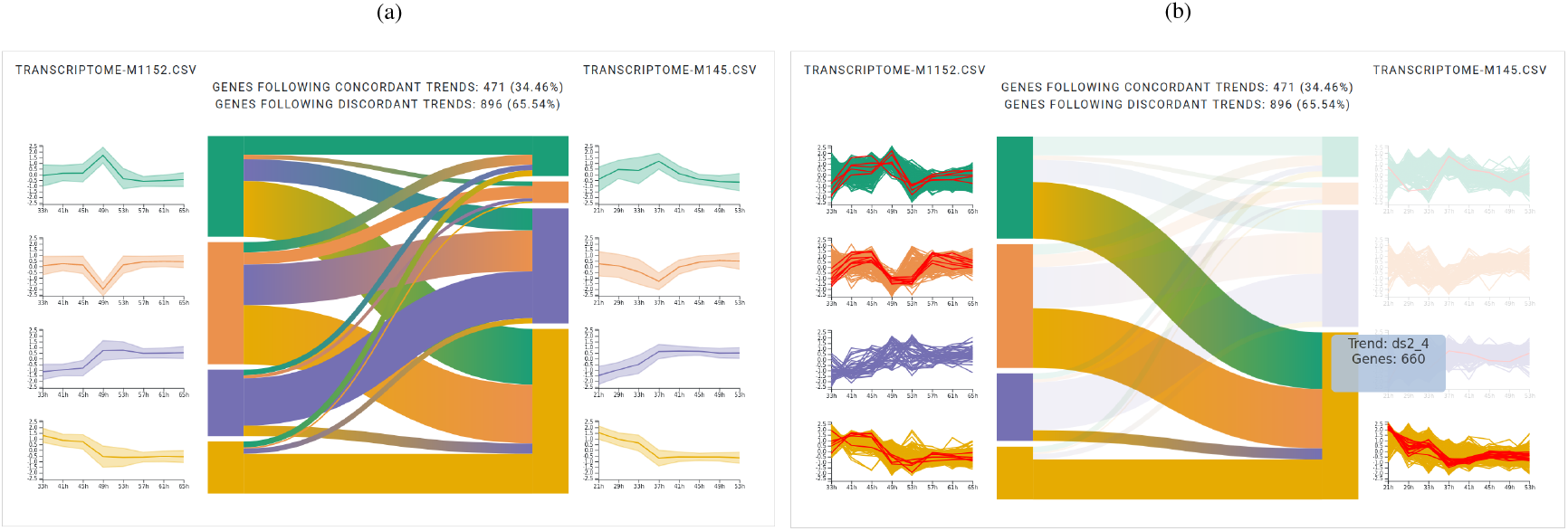
First-level analyses of the transcriptomes of the two *Streptomyces coelicolor* strains M1152 (on the left side in each subplot) and M145 (on the right side of each subplot) (a) after focusing on high-abundant genes in both data sets after removing genes with a median abundance below the 80th percentile. (b) A custom list containing genes involved in the metabolic switch to nitrate respiration under phosphate starvation has been uploaded and highlighted. The yellow node in the M145 is hovered.

These findings were investigated in more detail by combining the trend comparison with the information on genes known to be involved in the metabolic switch to nitrate respiration under phosphate depletion (Martín *et al*., 2017). The information on the corresponding gene IDs was uploaded to `OmicsTIDE` and the corresponding genes were highlighted in the profile diagrams (Figure 4b). Intriguingly, the majority of the these genes were detected in the green, orange and yellow cluster of the M1152 transcriptome and almost exclusively in the yellow cluster of the M145 transcriptome. The trend comparison revealed that the genes that follow the yellow trend in the wildtype strain, i.e. are down-regulated till timepoint 4, mainly follow the discordant green and orange trends, i.e. show a shifted alternating increasing and decreasing expression pattern in the mutant strain.

#### 5.2.2 *Inter-Omics, Intra-Condition*: M1152 transcriptome vs. M1152 proteome

The insights from the first trend comparison of the M1152 and M145 transcriptomes were used to further study the mechanisms in *Streptomyces coelicolor* by exploring how these findings are also manifested at the proteome level. Here, the pairwise trend comparison of the M1152 transcriptome and the M1152 proteome was used for this investigation. Especially the striking green and orange trends in the transcriptomes were found to show strong discordance across the two bacterial strains. Here, we exemplarily show for the green trend that this discordance is also reflected in the corresponding M1152 proteome (Figure 5). The hovering of the node in Sankey diagram that corresponds to the green trend in the M1152 transcriptome revealed 350 genes. Hereof only a small number of genes follows the concordant green trend in the M1152 proteome. However, the visual-interactive analysis allowed to detect that the remaining three trends in the proteome all share a small peak at a later time point. Since this small peak appears to a later time point than the peak in green trend of the M1152 transcriptome it could be used to further investigate whether this might suggest a time-delayed translation of the protein cognates corresponding to the gene expression in the green trend of M1152.

**Fig 5:**
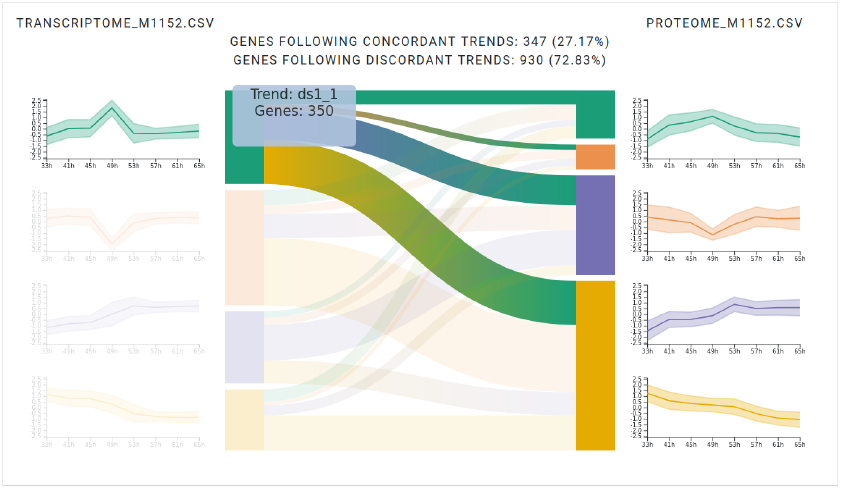
First-level analysis of M1152 transcriptome (left) vs. M1152 proteome (right) of the *Streptomyces coelicolor* data set with hovering on the green trend in the transcriptome. Only a small fraction of genes follows the same green trend in the proteome.

In summary, the parallel analysis of *intra-* and *inter-omics* data in `OmicsTIDE` leads to easily interpretable expression trends and possible hypotheses. Furthermore, the visual-interactive trend comparison makes even small changes in abundances salient to the user.

## 6 Discussion

In this paper we present `OmicsTIDE`, a new visualisation approach for the parallel exploration of multi-omics data. The parallel analysis of multi-omics data sets has become a powerful strategy to explore the informational flow propagating across different omics layers in a comprehensive manner. With `OmicsTIDE` we bridge the gap between separated single-omics and integrative multi-omics analyses. In the context of developing OmicsTIDE we also devised a classification system for multi-omics data, which offers an underlying framework for our tool, but may also serve useful for future developments in this field.

`OmicsTIDE` follows the strategy of a visual-interactive pairwise trend comparison. This strategy emphasizes the similarities and differences that appear at the interface of two omics data sets. With this, it marks an innovation compared to other tools that mainly aim to integrate a large number of omics data sets to derive a combined pattern. It should be noted that the pairwise analysis and the multi-omics integration are not mutually exclusive ways of analysis, but rather complement each other.

`OmicsTIDE` uses a Sankey diagram to compare trends across different data sets. With this visualization concordance and discordance in trends can be intuitively explored by directly linking the data sets. The trends themselves are either visualized using *centroid profile plots*, profile plots, or boxplots. While centroid profile plots visualize an overview of the profile, detailed profile plots show every gene separately. With this detailed visualization it is easier to track the behaviour of single genes. A magnified version of these plots can be found in the second level analysis, where it can be used to identify subclusters in a selection of genes. Profile plots are especially useful if the order of conditions is inherent (Gehlenborg, 2012). In contrast, boxplots do not assume that the conditions are ordered and are therefore better suited for categorical data. Moreover, they focus on visualizing the distribution of values at each condition. This is especially useful to identify outliers or for assessing data set consistency across replicates.

To compute trends from multi-omics data *OmicsTIDE* uses an *early integration* approach by first concatenating and then clustering the data. Currently, for the clustering `k-Means++` is implemented in `OmicsTIDE`. In addition the user can upload a custom clustering in `OmicsTIDE` and continue the visual exploration based on that.

The ability of `OmicsTIDE` to extract and compare trends was demonstrated in two case studies using different experimental designs. In the first case study, the integrated analysis of transcriptome and proteome data shows that `OmicsTIDE` can derive the most important information in few steps. The juxtaposition of the trends in each omics layer gave a clear overview of the distinct patterns in this data, while the interaction using hovering of the Sankey diagram confirmed the identification of the discordant and concordant dynamics between the two omics layers found by the authors of that respective study. These findings were further consolidated by a manual comparison of the genes extracted from the intersections of the trends in `OmicsTIDE` and the modules defined by the authors. Overall, we found that between 70 and 85% of the genes found in the respective modules agreed with the trends identified in `OmicsTIDE`.

The second case study applies a more complex experimental design enabling an *inter-omics* and *intra-omics* comparison. The central design feature of `OmicsTIDE` is the parallel trend comparison of two omics data sets. When more than two data sets are under investigation, as it was the case here, `OmicsTIDE` provides the option of combining different pairwise omics data comparisons within a single analysis. With this novel approach trends could be analyzed in the *inter-omics* as well as the *intra-omics* comparison while keeping an overview of all involved data sets. The combination of the different pairwise comparisons could reveal information that would not easily have been found by combining all of the used data sets in one integration. The exploration of the Sankey diagram using the *focus-on-hover* strategy could show that the trends initially found in the inter-condition analysis (the transcriptome comparison) are also revealed in the proteome.

## 7 Conclusion and Outlook

With `OmicsTIDE` we provide a visual analytics tool that is designed for biologists; its user interface creates clear default views that show the concordant and discordant patterns in omics abundance data in a pairwise manner. Even complex experimental designs, such as comparing intra-omics data as well as inter-omics data can be easily addressed in `OmicsTIDE`.

This tool marks an innovation in the pairwise comparison of data sets by reducing the information on the regulation of genes to single trends and allowing a clear visual-interactive comparison using the Sankey diagram. By following a multi-tab approach that separates the single analysis step, the switch between overview and the detailed view marks a striking difference to common interactive omics visualization tools.

While we were able to show that applying this approach extracts the main trends which can clearly be distinguished, we plan to implement more sophisticated clustering algorithms, such as `iCluster` (Shen *et al*., 2009). Such approaches might prevent biased trends, especially if the number of genes in one of the compared data sets is very high compared to the other data set. To counteract this bias in the current version of `OmicsTIDE` we analyze intersecting and non-intersecting genes separately, which guarantees an equal number of genes for both data sets in the intersecting analysis. Tools like `iCluster` would allow us to conduct a combined analysis for all genes.

For the determination of which genes commonly occur in the respective data sets, we compare the data sets through molecular IDs (e.g., gene or protein IDs), that greatly facilitates the comparative visualisation of the trends. The default view of `OmicsTIDE` focuses on the set of intersecting genes between the two data sets. Though also the trends of the non-intersecting genes can be visualised in profile plots, the current version of `OmicsTIDE` does not allow for a direct comparison. In a future version, a pairwise comparison could be achieved by categorizing the gene ID for example by common pathway IDs or other meta-information. An application could for instance be the investigation of orthologous genes in two different species. An example for this kind of analysis has been shown in a recent genome-wide comparative transcriptome analysis between *Arabidopsis thaliana* and *Zea mays* (Vercruysse *et al*., 2020), where trends of orthologous genes between these two species were compared.

## Acknowledgements

Funding for TH and JF was provided by the Deutsche Forschungsgemeinschaft (DFG, German Research Foundation)—Project-ID 398967434—TRR 261. MK was funded by the Carl-Zeiss-Stiftung.

